# Does ipsilateral remapping following hand loss impact motor control of the intact hand?

**DOI:** 10.1101/2023.04.19.537443

**Authors:** Raffaele Tucciarelli, Naveed Ejaz, Daan B. Wesselink, Vijay Kolli, Carl J. Hodgetts, Jörn Diedrichsen, Tamar Makin

## Abstract

What happens once a cortical territory becomes functionally redundant? We addressed brain and behavioural adaptations for the intact hand in individuals with a missing hand. Previous studies reported increased ipsilateral activity in the somatosensory territory of the missing hand (i.e., remapping) in acquired amputees, but not in individuals with a congenitally missing hand (one-handers). It is unclear whether remapping in amputees involves recruiting more neural resources to support the intact hand, and whether such activity is increased in tasks that demand greater motor control. We investigated sensorimotor learning and neural representation of the intact hand in one-handers and amputees using a multi-finger configuration task, as well as univariate and multivariate fMRI. We found that ipsilateral activity increased with motor demand - but only in the amputees group. However, these changes did not reflect behavioural differences. The representation of the finger configurations, as revealed by multivariate analysis, was stronger in amputees and closer to the typical representation found in controls’ contralateral hand territory, compared to one-handers. This collaborative contra-ipsilateral activity may reflect the intact hand’s efference copy. One-handers struggled to learn difficult finger configurations, but this did not translate to differences in univariate or multivariate activity relative to controls. Together with a supplementary structural white matter analysis, our results suggest that enhanced activity in the missing hand territory may not reflect intact hand function. Instead, we suggest that plasticity is more restricted than generally assumed and may depend on the availability of homologous pathways acquired early in life.

**Significant Statement:** We studied whether brain resources in the missing-hand territory support demanding intact hand motor control in people who were born with one hand or lost a hand later in life. We found that amputees had increased activity in the brain area used for the missing hand, but no improvement in the performance of their intact hand. This collaborative contra-ipsilateral activity may reflect the intact hand’s efference copy. One-handers showed slight deficits while learning to perform complex motor movements, but no brain activity differences in the missing hand territory, compared to controls. Our results suggest that brain plasticity is limited and may depend on early life experiences.

## Introduction

Specific functions of mature cortical areas are determined by their molecular properties, their histological organization, and intrinsic and extrinsic connectional fingerprints. The unique identity of a given area is determined by genetic expression and is moderated by electrical activity over the course of early development (see Sur and Rubenstein, 2005 for review). This phase of increased susceptibility to input in shaping the neural circuit is called a *critical period* (Levelt and Ḧubener, 2012). The critical period might be enabled because of plasticity ‘brakes’, such as inhibitory circuits, neural over-growth and synaptic pruning, normally affording homeostatic balance, have not yet been finalised (Takesian and Hensch, 2013). Yet, even in these earliest stages of development, it seems that the assignment of brain function to a given cortical structure is largely fixed. For example, fMRI studies of children who sustained left hemisphere perinatal stroke found that the (typically left dominant) language areas in the inferior frontal cortex were located in anatomically homologous areas in the right hemisphere (Tillema et al., 2008; Raja Beharelle et al., 2010; Tuckute et al., 2022). In this context, it is interesting to consider how a redundant cortical area’s function changes after hand loss, either because of congenital hand malformation or acquired arm amputation later in life.

If brain plasticity is more potent in early development, we might expect to find more functional changes in individuals born with a missing hand due to a developmental malformation (hereafter – one-handers), in comparison to individuals who only lost their hand later in life (hereafter – amputees). In particular, the intact hand should primarily benefit from the redundant resources in the missing hand territory to adapt better to life with only one hand. If this reallocation of resources from the missing hand territory towards the intact hand is functional, we might expect to see improved motor abilities and learning in one-handers relative to controls. Surprisingly, previous research reported increased activity in the missing hand area territory for inputs and outputs from the (ipsilateral) intact hand in amputees but not one-handers (Kew et al., 1994; Hamzei et al., 2001; Bogdanov et al., 2012; Makin et al., 2013a; Philip and Frey, 2014; Valyear et al., 2020). While one-handers show profound alterations to the classical sensorimotor ‘homunculus’, where the deprived hand area was demonstrated to be activated by multiple body parts (e.g. arm, face, feet, torso), it does not appear to be activated by the intact hand (Hahamy et al., 2017; Hahamy and Makin, 2019). It has therefore been suggested that the brain may first need to establish a functional connection between the two hands through bimanual experience to enable ipsilateral functionality (Philip et al., 2015). Another possibility is that the tasks used in previous studies were not sufficiently challenging (e.g., opening and closing the hand), and therefore did not require recruitment of additional processing. In other words, if ipsilateral processing due to plasticity processes provides *additional* resources to aid motor control of the intact hand, it may require difficult tasks to activate it.

Here we aimed to address the relationship between brain and behavioural adaptations for the intact hand in individuals with a missing hand. To better gauge whether activity changes in one-handers and amputees are functional, we varied task difficulty. Participants had to simultaneously press with three fingers onto a piano-like keyboard, while keeping the other two fingers relaxed. Some combination of fingers are known to be more difficult than others (Waters-Metenier et al., 2014), and we therefore selected two sets of 5 configurations that systematically ranged from easy to difficult. Motor difficulty is known to increase activity level, particularly in the ipsilateral hemisphere (Verstynen et al., 2005). Therefore, we used brain scans while participants performed the same task to compare the net activity (remapping) and multivariate representational similarity (information content and representational structure) across the primary somatosensory and motor territories (S1 and M1) of the missing and intact hand. We focused on S1 as it is known to contain greater information content relating to finger movements (Schieber, 2001; Ejaz et al., 2015). To explore the structural underpinnings of these functional changes, we also used diffusion MRI to examine potential white matter microstructural changes in transcallosal fibre connections linking the two hand areas.

We predicted that, if one-handers rely on ipsilateral processing for difficult tasks, we should see increased activity and information content in the missing hand cortex compared to controls, leading to improved performance compared to controls. Alternatively, if the functional availability of homologous resources depends on bimanual experience, we should expect to find greater activity and information content in the missing hand territory of amputees relative to controls. Moreover, this information should be organised in a homologous representational structure relative to the intact hand territory.

## Materials and Methods

The experimental procedures described in this manuscript were run as part of a larger study (the full study protocol can be found on https://osf.io/gmvua/). Here we focus on procedures related to the finger coordination task. The motor task was similar to previous studies (Waters-Metenier et al., 2014; Ejaz et al., 2015). Participants took part in the training session outside the scanner first and then an fMRI session.

### Participants

Amputees (N=19; Mean Age=49.05±12.05), one-handers (N=16; Mean Age=43.44±11.40), and two-handed controls (N=16; Mean Age=45.37±10.67) were invited to take part in a motor control task. The current experiment was comprised of a training session outside the scanner, followed by a scanning session. Not all participants were able to take part in (or complete) the scanning session. Thus, the final sample in the scanning session was N=16 amputees (Mean Age=48.40±12.6), N=13 one-handers (Mean Age=45.80±11.30), and N=14 two-handed controls (Mean Age=44.20±12.20). Furthermore, due to technical reasons, we could not register the responses of two amputee participants in the scanner, therefore the analyses of the finger coordination task during the MRI session were based on N=14 amputees (Mean Age=49.5±13.1). For the DTI, we were able to collect data for N=18 amputees, N=13 one-handers, and N=13 controls. The mean age was not significantly different between groups (Motor training: F_(2,48)_=1.10, p=.341, η_p_=.034; Scanner: F_(2,39)_=1.11, p=.340, η_p_=.054). Nevertheless, to take into account any potential inter-individual impact of age, we included participants age as a covariate in the analyses. In all analyses, outliers were defined as values exceeding the metrics of interest of three standard deviations from the mean. For the behavioural data during the training session, we identified one potential outlier, but since this outlier did not impact the results qualitatively, we opted to include the outlier in the final analysis. For the multivariate data, we identified an outlier that we decided to remove because its removal qualitatively changed the significance level.

Recruitment was carried out in accordance with the University of Oxford’s Medical Sciences inter-divisional research ethics committee (MS-IDREC-C2-2015-012). Informed consent and consent to publish was obtained in accordance with ethical standards set out by the Declaration of Helsinki. All participants were compatible with local magnetic resonance imaging (MRI) safety guidelines.

### Apparatus

Responses were recorded using a custom-built 5-finger MRI-compatible piano-like device (Wiestler and Diedrichsen, 2013; Ejaz et al., 2015; Wesselink et al., 2019). Each key was equipped with a sensor that could continuously measure isometric force during a finger press. The sensors were connected to a laptop and the applied forces were monitored online. Participants received real-time visual feedback on how much force each finger exerted by means of moving horizontal white cursors corresponding to each key. In the training task outside the scanner, the apparatus was placed on a desk in front of the seated participant, who rested the five fingers of their intact hand (or dominant hand in controls) on the keys that were immobile but able to measure the applied pressure. Participants could choose to keep their fingers extended or flexed, based on comfort. Inside the scanner, the device was placed on their lap or belly, depending on their preference.

### General procedure

*Instructions:* The top of the screen showed five vertical grey bars, each corresponding to one of the keys. At rest, participants were required to apply and maintain a minimal force (0.5N) on the keys, as indicated by a horizontal bar at the bottom of the screen (hereafter baseline area). In a typical trial, three of the vertical bars turned green indicating which of the keys to press. Participants were instructed to wait until the appearance of a go cue that was provided as a green horizontal bar similar in dimension and right above the baseline area. At this point, participants had to press three keys in synchrony (chord-like configuration) and using the same force (2.5N) on all instructed fingers while keeping the non-instructed fingers placed relaxed on the keys. In this way, participants had to use the sensory information provided by all fingers which is fundamental in dexterous manipulation (Pruszynski et al., 2016). Participants received a positive feedback (i.e., a point) as soon as they configured the instructed fingers as required. Once the finger cursors were successfully stabilised in the target area, the area disappeared indicating the participants to go back to the baseline position (this was a requirement to obtaining a point in the next trial) by releasing the pressure on the instructed fingers. At this point, a new trial started. Note that the training session was self-paced, whereas the scanning session was timed (see below for details).

*Training session:* First, the experimenter explained the task and showed the participants how to use the device. Then, the participants performed a few familiarization trials with a set of configurations not used in the study. This was followed by a single-finger movement block, where the participants had to press only one of the five fingers. This bock was repeated one more time at the end of the training (as detailed in Wesselink et al., 2019). Then, the actual training session started and it lasted 25 minutes. Within this time window, participants were encouraged to complete as many blocks as possible. Each block was about 3 minutes-long, depending on the performance, leading to a variable number of blocks across participants. On average, participants completed M_all_=5.69 blocks (SD_all_=1.42; M_Amputees_= 5.58, SD_Amputees_= 1.54; M_One-handers_= 5.62, SD_One-handers_= 1.36; M_Controls_= 5.87, SD_Controls_= 1.41), and the three groups did not differ for the number of blocks completed (F_(2,48)_=0.20, p=.817, η_p_=.01). Visual instructions of the required chord were presented for 3 seconds, followed by the go cue. Within each block, instructions for the same finger configuration were repeated 4 times.

*fMRI session:* After the training session, participants were invited to take part in a similar motor task as part of the fMRI study. In the scanner, there was no minimal pressure requirement at baseline because the application of constant pressure was tiring while lying supine. In addition, the task was timed. Visual instructions were presented for 1.3 seconds, and participants had to execute the chord (i.e., press and release the keys) within 2.3 seconds from the onset of the instructions. The same instruction was repeated three time resulting in blocks of 6.9 seconds. Each finger configuration block was repeated three times within a run, resulting in 45 trials per run (9 trials by 5 finger configurations). Participant took part in four runs and each run lasted around 3.5 minutes (141 volumes).

*Behavioral performance*: Behavioural performance was measured as the deviance from the required finger configuration, by taking into account two sources of error: 1) any deviation of the non-instructed fingers from the baseline (0.5 N); and 2) any deviation of the each instructed fingers from the average force as all the instructed fingers were expected to exert a similar force (2.5 N). These two forms of residuals were computed within the response and release time, summed up and averaged across time to obtain a unique measure of performance per trial (see Waters-Metenier et al., 2014). In line with previous studies (Ejaz et al., 2015; Waters et al., 2017), the beginning of the response was defined as the point in time in which at least one of the fingers exceeded the threshold of 1.5 N when pressed.

### Finger configuration and difficulty levels

Figure 1 displays the configurations used in the training session (Panel A, easy to difficult from the bottom to the top) and in the scanner (Panel D, easy to difficult from the bottom to the top). In the training session, we used the following finger configurations (1: Thumb, 2: Index, 3: Middle, 4: Ring, 5: Little finger): 345, 123, 124, 245, 135. The aim of the training session was twofold: 1) measure the sensorimotor learning of participants, and 2) familiarise participants with the task before entering the scanner. In the scanner, we used different finger configurations (145, 234, 134, 125, and 235) in order to minimise any differences across groups that were hypothesised to arise due to different training capacity.

**Figure 1.**
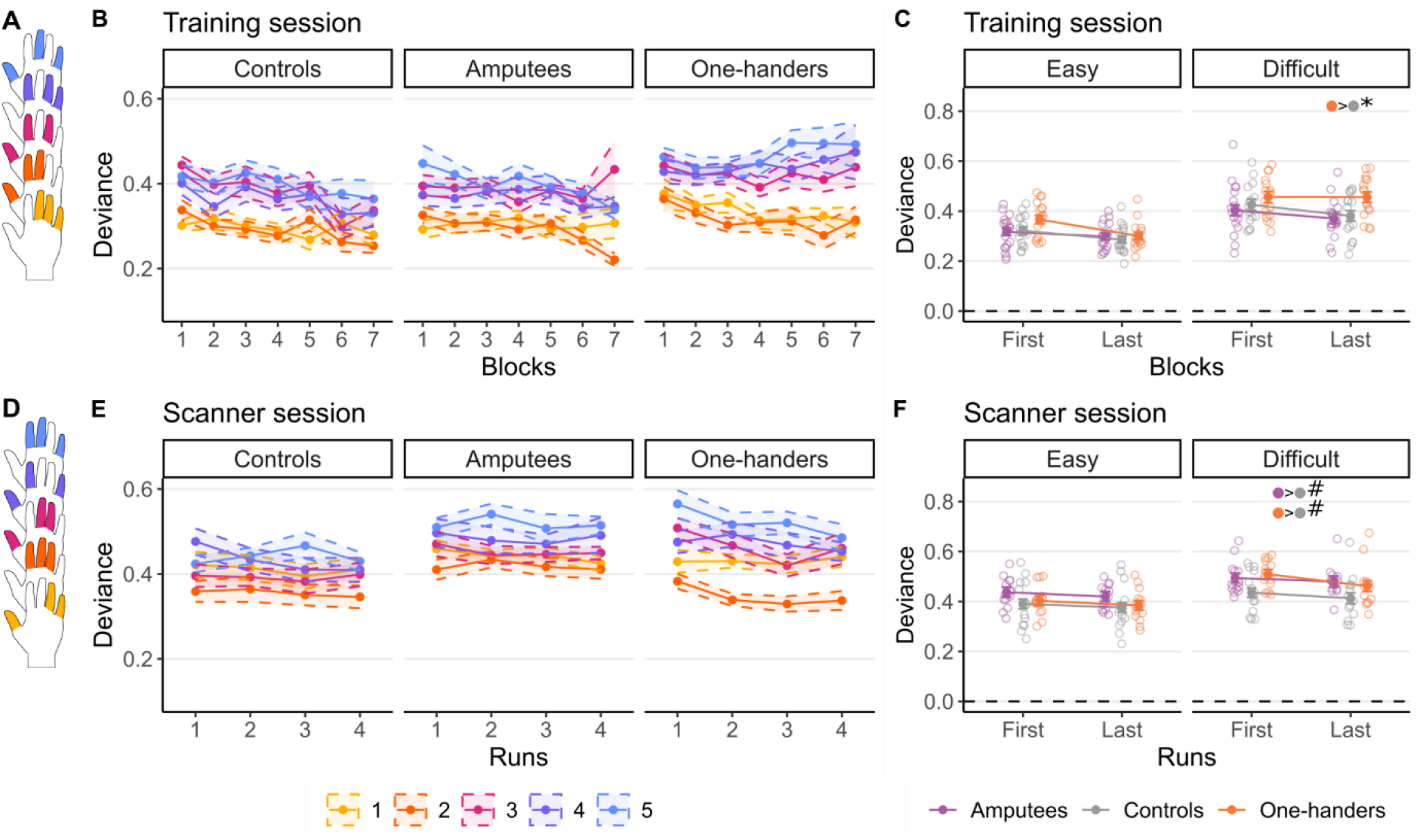
Intact hand motor performance. Schematic representation of the finger configurations used for the motor task during the **(A)** training and **(D)** fMRI sessions. The colours represent the graded difficulty across configurations (based on the inter-finger enslavement components), with colder colours indicating more difficult configurations. **(B, E)** Line plots of the mean deviance values (± SEM) across blocks/runs for the (B) training and (D) fMRI sessions. Deviance scores reflect the extent to which the pressure exerted by the 5 fingers deviated from the expected configuration (see *General procedure* in the method section). Smaller deviance reflects better performance. **(C, F)** Effect plots showing the marginal means of the deviation values predicted by the model (repeated measures ANCOVA, controlling for age) for the (C) training and (F) fMRI session, as well as individual participants performance (grey dots). For the training session (C), deviance was averaged over the easy configurations (345, 123, and 124) and difficult configurations (245 and 135), for the first and last blocks each participant performed. For the fMRI session (F), deviance was averaged over the easy (145 and 234) and difficult (134, 125, and 235) configurations, for the first and 4th run. In the training session, participants showed improvement in motor control, as indicated by a decrease in deviance, but one-handers demonstrated reduced learning for the most challenging configurations. In the fMRI session, both one-handers and controls showed reduced performance, relative to controls.

To independently confirm the previously estimated difficulty levels of finger configurations, defined from a pilot session of a previous study (Waters-Metenier et al., 2014), were appropriately labelled, we also utilized a model-based approach. To this aim, we used the amount of flexion enslavement (% of maximal voluntary contraction) between fingers in a single-finger task (Yu et al., 2010). In particular, we reasoned that easy configurations would be characterized by high amount of enslavement between the instructed fingers, high amount of enslavement between the non-instructed fingers, and low amount of enslavement between the instructed and non-instructed fingers. For each chord, we estimated three components of enslavement: the total (i.e., sum) of enslavement for the instructed fingers (E1), the enslavement for the non-instructed fingers (E2), and the enslavement between the instructed and non-instructed fingers (E3). Then, we combined the three components (i.e., E1+E2-E3) to obtain a unique measure of enslavement such that the configurations with a high score were categorized as easier than the ones with a low score. Using this measure, we sorted the configurations from easy to difficult and divided them into two groups: *easy* (345, 145, 234, 123, 124) and *difficult* (134, 125, 245, 235, 135). In our analysis, for the training session, the *easy* averaged configurations were 345, 123, and 124, and the *difficult* averaged configurations were 245 and 135; for the scanning session, the *easy* averaged configurations were 145 and 234, and the *difficult* averaged configurations were 134, 125, and 235. We also used this scoring to establish the easiest and most difficult configurations for specific fMRI analysis.

### MRI data acquisition

MRI images were acquired using a 3T MAGNETON Prisma MRI scanner (Siemens, Erlangen, Germany) with a 32-channel head coil. Functional images were collected using a multiband T2*-weighted pulse sequence with a between-slice acceleration factor of 4 and no in-slice acceleration (2 mm isotropic, TR: 1500 ms), covering the entire brain. The following acquisition parameters were used: TE: 32.40 ms; flip angle: 75°, 72 transversal slices. Field maps were acquired for field unwarping. A T1-weighted sequence was used to acquire an anatomical image (TR: 1900 ms, TE: 3.97 ms, flip angle: 8°, spatial resolution: 1 mm isotropic). Diffusion-weighted MRI (dMRI) data were acquired using the following parameters: TR: 2951 ms, TE: 79.80 ms, flip angle: 80°, spatial resolution: 1.5 mm isotopic, 84 transversal slices. Gradients were applied along 60 uniformly distributed directions with a b-value of 1000 s/mm2. Five non–diffusion-weighted images with b = 0 s/mm2 were also acquired. No task was given to the participants during the structural and DTI acquisition. They viewed a calm nature video to prevent them from falling asleep and making large head movements.

### fMRI preprocessing and first-level analysis

MRI data were preprocessed using a standard pipeline as implemented in FSL 6 (Smith et al., 2004; Jenkinson et al., 2012). The following steps were applied to each functional run: motion correction using MCFLIRT (Jenkinson et al., 2002); B0 fieldmap correction to account from distortions due to magnetic field inhomogeneity; brain extraction using BET (Jenkinson et al., 2002); high-pass temporal filtering of 90 s; and spatial smoothing with a Gaussian kernel of full with at half maximum (FWHM) of 5 mm for the univariate analyses and 3 mm for the multivariate analyses.

In order to estimate brain activity related to our configuration task, we employed a voxel-based general linear model (GLM) as implemented in FEAT. For each functional run, time series were predicted using five regressors of interest corresponding to the five configurations that participants had to do in the scanner. These regressors were convolved with a double-gamma function and their temporal regressors were also added to the design matrix to account for temporal variability of the BOLD response. We also included the motion parameters resulting from the MCFLIRT step, and columns indicating outlier volumes as returned from the FSL function *fsl_motion_outliers* with default and recommended parameters (root mean square intensity difference of each volume to the reference volume as metric; as a threshold, metrics that were larger than 75^th^ percentile+1.5*InterQuartile rage were considered outliers). The number of volume outliers was small for all groups (Amputee group: mean proportion volumes excluded= 0.044±0.012; One-handers group: mean proportion volumes excluded= 0.045± 0.017; Control group: mean proportion volumes excluded= 0.047±0.019), and there was no difference between the three groups (F_(2,40)_= 0.152, p=.860, η_p_=.008).

### MRI analysis

For each individual, cortical surfaces were estimated from the structural images using Freesurfer 5.3.0 (Dale et al., 1999; Fischl et al., 2001). Further MRI analyses were also implemented using Connectome Workbench software (https://www.humanconnectome.org/software/connectome-workbench). To define the ROIs, we used the Brodmann Area (BA) maps included in Freesurfer (Fischl et al., 2008) that are based on the histological analysis of ten human post-mortem brains.

### ROI definition

We focused our analyses on bilateral hand S1 (and area BA3b in particular), which has been most commonly associated with remapping in animal and human (see Makin and Flor, 2020 for a literature overview) studies. Further motivation for our S1 focus is that previous research has consistently shown that S1 contains more finger information (including inter-finger configurations) relative to M1 (Schieber and Hibbard, 1993; Ejaz et al., 2015). This is because S1 topography tends to be well-defined, relative to M1 where the information content is more widespread (Schieber, 2001; Graziano and Aflalo, 2007). However, we also report results from M1 (area BA4). The ROIs were defined in the fsaverage template space using probabilistic cytoarchitectonic maps (Fischl et al., 2008), based on 2.5 cm proximal/distal (Wiestler and Diedrichsen, 2013; Berlot et al., 2019; Ogawa et al., 2019; Arbuckle et al., 2022) to the hand knob (Yousry et al., 1997). The resulting hand S1 was then projected to the individual reconstructed surfaces. Here we focused on nodes with at least 50% probability of being part of BA3b. We chose this threshold to make sure that all of BA3b was included, and to make sure the regions were large enough. However, we note that given the inherent smoothness of the data, our preprocessing procedure and the probabilistic nature of the anatomical atlas, the ROIs are likely to contain relevant activity from neighbouring S1 areas. We then mapped the surface ROIs to the individual volumetric high-resolution anatomy and resampled to the lower resolution functional brain. Hand M1 was defined in a similar way as hand S1 described above. As a control region, we used hMT+ that was created combining area FST, V4t, MT and MST from the Human Connectome Project parcellation (Glasser et al., 2016).

### Representational similarity analysis (RSA)

Information content within each ROI was estimated using RSA (Kriegeskorte et al., 2008). For each participant and run, we extracted the first-level betas estimated with FEAT (see previous section *fMRI preprocessing and first-level analysis*) from each ROI and computed the pairwise cross-validated Mahalanobis (or crossnobis) distance (Walther et al., 2016) between chord-related beta patterns as a measure of their dissimilarity. Multidimensional noise normalisation was used to increase reliability of distance estimates (noisier voxels are down-weighted), based on the voxel’s covariance matrix calculated from the GLM residuals. The advantage of using the crossnobis distance is twofold: 1) spatially correlated noise is removed using multivariate noise normalization and this improves the estimate of the dissimilarities (Walther et al., 2016); 2) cross-validation ensure that if two patterns only differ by noise, their mean dissimilarity estimate will be zero. As a consequence, the dissimilarity between two patterns can also be negative (Diedrichsen et al., 2016) and thus dissimilarities significantly larger than zero can be taken as evidence that the two patterns are distinct and that the ROI contain task-related information (e.g., distinct representation of configurations). The crossnobis dissimilarity was computed using the python library for RSA *rsatoolbox* version 0.0.4 (https://github.com/rsagroup/rsatoolbox).

### Diffusion MRI preprocessing

Diffusion data were preprocessed using a custom pipeline that combined tools from MRItrix 3.0 (Tournier et al., 2019), ExploreDTI 4.8.6 (Leemans et al., 2009), and FSL 5.0.9 (Smith et al., 2004; Jenkinson et al., 2012). These included: 1) de-noising using the MP-PCA (principal component analysis of Marchenko-Pastur) method in MRtrix (Veraart et al., 2016); 2) Gibbs ringing correction using ‘mrdegibbs’ in MRItrix (partial Fourier; Kellner et al., 2016); 3) Global signal drift correction using ExploreDTI (Vos et al., 2017); and 4) Motion EPI distortion correction using Eddy and Topup within FSL (Andersson et al., 2003). Data were visually checked as part of quality assurance procedures. Whole brain voxel-wise maps of fractional anisotropy (FA) and mean diffusivity (MD) maps were then derived from the preprocessed data by fitting the diffusion tensor model. FA represents the degree to which diffusion is constrained in a particular direction, and ranges from 0 (isotropic diffusion) to 1 (anisotropic diffusion). MD (10^-3^mm^2^s^-1^) represents the average diffusivity rate. The diffusion tensor was estimated and fitted using the nonlinear least squares method with Robust Estimation of Tensors by Outlier Rejection (RESTORE) applied (Chang et al., 2005).

### Tractography

A multiple-ROI tractography approach enabled specific transcallosal pathways to be constructed in each participant between their left and right S1 hand areas (see also Postans et al., 2020). Initially, each participant’s ROIs in T1 space (see Section *ROI definition* above) were registered to their native space diffusion MRI image using the following steps: 1) the T1-to-diffusion transformation matrix was generated using FLIRT with 6 degrees-of-freedom and the correlation ratio cost function. The fractional anisotropy (FA) map was used as the reference image (rather than the b0 image) as it provided better image contrast; 2) the transformation matrix was then applied to the individual subject ROIs in T1 space using FLIRT. As tractography can be challenging from grey matter ROIs (due to low anisotropy), the diffusion space ROIs were dilated by 1.5 mm to include some white matter voxels (Thomas et al., 2014).

Tractography was initially performed from all voxels in the left hemisphere ROI in each participant’s native diffusion MRI space in ExploreDTI (v4.8.3; Leemans et al. 2009) using a deterministic tractography algorithm based on constrained spherical deconvolution (Tournier et al. 2008; Jeurissen et al. 2011). Spherical deconvolution approaches enable multiple peaks to be extracted in the fibre orientation density function within a given voxel, allowing complex fibre arrangements, such as crossing/kissing fibres, to be modelled more accurately (Dell’Acqua and Tournier, 2019). The contralateral S1 ROI was then used as an “AND” gate to capture any streamlines that arose from the seed ROI and terminated in the contralateral ROI. Next, the same procedure was repeated, this time starting with the right hemisphere ROI as seed and gating with the right hemisphere. This process was conducted for each participant and then inspected visually by the research team (CJH, RT). A step size of 0.1 mm and an angle threshold of 60° were applied to prevent the reconstruction of anatomically implausible streamlines. Tracking was performed with a supersampling factor of 4 × 4 × 4 (i.e., streamlines were initiated from 64 grid points, uniformly distributed within each voxel). The resulting inter-hemispheric pathways were then intersected with the whole-brain voxel-wise FA and MD maps (see above) to derive four tract-specific measures of microstructure in each participant (S1-to-S1 and M1-to-M1, in both directions). As in Postans et al. (2020), the FA and MD values for the left-to-right and right-to-left segments were combined into a streamline-weighted mean using the following equation:

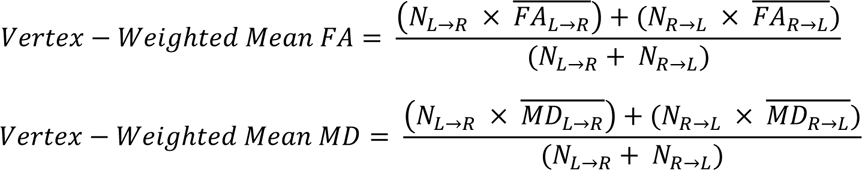

### Tract-based spatial statistics (TBSS)

We also conducted a complementary voxel-wise statistical analysis of the FA and MD data using TBSS (Smith et al., 2006). First, each participant’s FA and MD maps were aligned to the standard MNI template using nonlinear registration (Andersson et al., 2010). Second, the mean FA image was created and subsequently thinned (using the default FA threshold = 0.2) to generate the mean FA skeleton, which represents the centre of all tracts common to the group. Third, participants’ FA and MD data were projected onto the skeleton for voxel-wise analyses using randomise in FSL (Winkler et al., 2014). For both FA and MD, a general linear model was constructed, which specified contrasts between amputees and one-handers (amputees > one-handers, and one-handers > amputees), and also each experimental group against controls. Age (de-meaned) was added as a covariate. Following prior work *(Hahamy et al., 2015)*, analyses were first restricted to the bilateral corticospinal tract using an ROI mask [labelled “WM Corticospinal tract”] from the Julich Histological Atlas *(Amunts et al., 2020)* using threshold free cluster enhancement (TFCE) with a corrected alpha of 0.05. We also conducted an additional whole brain analysis to examine any potential group difference outside our main ROIs (using the same TFCE-corrected threshold of p = 0.05). All reported TBSS co-ordinates are in MNI 152 space.

### Statistical analysis

Statistical analyses were performed using custom-made scripts written in Matlab R2020b (The MathWorks), R version 4.1.3 (R Core Team, 2022) with RStudio (2021.09.0 Build 351), python 3.10.6 with spyder 5.3.3, and JASP 0.17. Behavioural performance (mean deviation) for the training and the scanning sessions were analysed using three-way repeated-measures ANCOVAs (rmANCOVAs) with age (de-meaned) included as a covariate, group as a between-subject factor, and block number and difficulty as within-subject factors. Brain activity (z scores, averaged across runs) for each ROI was analysed using a three-way rmANCOVA with age included as a covariate, group as a between-subject factor, and hemisphere and difficulty as within-subject factors. To test for existing information content, dissimilarities were tested against zeros using a two-tailed one-sample t-test for each group and hemisphere. Dissimilarities were also analysed in two ways. In one analysis, we only selected the easiest and most difficult finger configuration pairs and used a three-way rmANCOVA with age included as a covariate, group as a between-subject factor, and hemisphere and difficulty as within-subject factors. In a second analysis, we averaged across all finger configuration pairs and ran a two-way rmANCOVA with age included as a covariate, group as a between-subject factor, and hemisphere as a within-subject factor. To test for existing functional homotopy (i.e., correlation between finger configuration pairs across hemispheres), we used two-tailed one-sample t-test for each group and hemisphere. We also used a one-way ANCOVA with age as a covariate and group as a between-subject factor to investigate differences in functional homotopy between groups. To investigate similarity to typical contralateral representation in the experimental groups (i.e., correlation between the RDMs of the experimental participants, amputees and one-handers, with the contralateral RDM averaged across the control participants), we used two-tailed one-sample t-test for each group and hemisphere. We also used a two-way rmANCOVA with age as a covariate, group (one-handers, amputees) as a between-subject factor, and hemisphere as a within-subject factor to investigate differences in typical contralateral representation between the experimental groups. Prior to these analyses, correlation values were standardized using the Fisher’s r-to-z transformation. Independent t-tests were used to test for group differences. The experimental groups (amputees and one-handers) were compared against the control group unless differently specified. To control for age while performing an independent t-test, we first ran an ANCOVA and then computed the contrasts of interest using the R package *emmeans 1.8.2*. For post-hoc comparisons that were exploratory (that is, not a priori and not confirmatory), we adjusted our significance alpha level for multiple-comparisons using the Bonferroni approach. In the results section, we report the uncorrected p-values with a note of the adjusted alpha level. For non-significant results of interest, we reported the corresponding Bayes Factor (*BF_10_*), defined as the relative support for the alternative hypothesis. While it is generally agreed that it is difficult to establish a cut-off for what consists sufficient evidence, we used the threshold of *BF*<1/3 as sufficient evidence in support of the null, consistent with others in the field (Wetzels et al., 2011; Dienes, 2014). For Bayesian ANCOVAs, we used a uniform model as a prior and for Bayesian t-tests, we used the Cauchy model with a width of 0.707, which are the default settings in JASP. For all analyses, whenever the normality assumptions were not met, we adopted a permutation approach using the function *aovperm* of the R package *permuco 1.1.2* with default settings (permutation method for fixed effects models: *freedman_lane*; for mixed effects models: *Rd_kheradPajouh_renaud*) and we report these results with a note only when they are qualitatively different from the parametric approach.

### Data code and accessibility

The preprocessed data and the scripts necessary to reproduce the analyses can be found at https://osf.io/hsvkc/.

## Results

### One-handers show reduced benefits from brief training of difficult finger configurations

We first explored whether individuals with a missing hand, either due to congenital malformation (one-handers) or amputation in adulthood (amputees), differ from controls in their ability to learn to perform a finger configuration task with varying levels of difficulty. Mean deviations from the instructed hand configuration for each of the 5 configurations across the first 7 blocks are shown for the three groups in Figure 1B, with more difficult configurations displayed in cooler colours. At the first attempt (block 1), there was no difference in performance between the experimental and control groups, except for a trend for the most difficult level, in which one-handers showed worse performance compared to the control group (10 comparisons, no corrections for multiple comparisons). To quantify training effects across groups, we averaged deviation means between easy (configurations 1-2) and difficult levels (configurations 3-5) for each participant, and compared performance between the first and last blocks completed during training (Figure 1C). The resulting 3 (group) x 2 (block) x 2 (difficulty) ANCOVA (controlling for age) resulted in a significant 3-way interaction (F_(2,47)_=5.28, **p=.009**, η_p_=0.18), indicating that participants across the 3 groups benefited differently from the practice, with respect to difficulty levels. In addition, a main effect of difficulty (F_(1,47)_=137.22, **p=<.001**, η_p_=0.74) and block number (F_(1,47)_=17.48, **p=<.001**, η_p_=0.27) was found, with no significant main effect of group (F_(2,47)_=1.31, p=.280, η_p_=0.05). As apparent from the figures, this was driven by a lack of learning effect in the one-handed group, specifically for the difficult configurations. To better quantify this, we ran a separate repeated-measures ANCOVA for each group and observed a significant interaction between block number and difficulty for the one-handers only (F_(1,14)_=12.61, **p=.003**, η_p_=0.47). To further explore the differential learning effect observed in the one-handed group, we compared differences in performance between the last and the first block (Figure 1C), and found a significant learning effect in the easy condition only (Easy: t_(14)_=-3.58, **p=.003**; Difficult: t_(15)_=.14, p=.889, BF_10_=0.258; Bonferroni adjusted α: .05/2=.025), suggesting that the impairment in learning was specific for the difficult configurations. This was also confirmed by significant differences in the last block of training between one-handers and controls for the difficult configurations only (t_(47)_=2.32, **p=.024**).

We next examined whether these group differences in performance were replicated in the fMRI task, where 5 different configurations were used Figure 1D. Figure 1E shows performance across the 4 runs. To test for differential learning effects, we repeated the analysis mentioned above while comparing performance across groups and difficulty levels between the first and the last runs (Figure 1F). The 3-way interaction in the ANCOVA was not significant (F_(2,38)_=.48, p=.622, η_p_=0.02), indicating that the groups did not show different learning effects – indeed as shown in the figure, performance had already plateaued. However, we did observe a significant interaction between group and difficulty (F_(2,38)_=3.93, **p=.028**, η_p_=0.17), indicating that participants in different groups responded differently to task difficulty. We also observed a trend towards a main effect of group (F_(2,38)_=2.71, p=.080, η_p_=0.12), in addition to a main effect of difficulty (F_(1,38)_=73.61, **p=<.001**, η_p_=0.66) and block number (F_(1,38)_=9.89, **p=.003**, η_p_=0.21). The interaction between group and difficulty only showed a trending result and was driven by the one-handers performing worse on the difficult configurations relative to controls (t_(38)_=2.38, p=.022, Bonferroni adjusted α: .05/3=.0167). This is reflective of the behavioural results found outside the scanner, where the one-handers showed worst performance on the difficult configurations at the end of the training session. Here, we also found a trending result suggesting performance deficits in the amputee group relative to controls in the difficult configurations (t_(38)_=2.42, p=.021, Bonferroni adjusted α: .05/3=.0167). However, the one-hander and amputees groups did not differ relative to each other in performance (t_(38)_=0.06, p=.955, Bonferroni adjusted α: .05/3=.0167). It is important to note that previous tests comparing the two experimental groups against the control group only showed a trend (i.e., did not survive the multiple comparisons correction as the Bonferroni corrected p-values were both below .067), and as such, these findings should be interpreted with caution.

### Amputees show increased averaged ipsilateral activity that scales with difficulty

Next, we examined univariate activity levels across the bilateral S1 hand regions of interest (ROIs, Figure 2A). To estimate whether difficulty modulated brain activity differently for the different groups and hemispheres, we first conducted a 3-level ANCOVA, including 3 (group) x 2 (hemisphere) x 2 (difficulty) and age (as a covariate). We observed a significant 3-way interaction (F_(2,39)_=5.23, **p=.010**, η_p_=0.21), confirming that difficulty modulates activity differently across hemispheres and groups, as shown in Figure 2B. The analysis also revealed a significant interaction between group and hemisphere (F_(2,39)_=6.46, **p=.004**, η_p_=0.25), difficulty and hemisphere (F_(1,39)_=6.16, **p=.018**, η_p_=0.14), and main effects of hemisphere (F_(1,39)_=71.02, **p=<.001**, η_p_=0.65) and difficulty (F_(1,39)_=11.33, **p=.002**, η_p_=0.23). To further explore the 3-way interaction, we conducted two separate 2-level ANCOVAs for each hemisphere. As hypothesised, we observed group differences within the ipsilateral cortex only, where we found a significant interaction between group and difficulty (F_(2,39)_=3.39, **p=.044**, η_p_=0.15) and a main effect of group (F_(2,39)_=5.95, **p=.006**, η_p_=0.23), while no main effect or interaction involving group was observed in the contralateral hemisphere (all p>.6). This suggests that activity scales with task difficulty differently across groups in the ipsilateral cortex (which is the missing hand cortex in the experimental groups). The main effect of difficulty was significant in both hemispheres (Contralateral: F_(1,39)_=15.68, **p<.001**, η_p_=0.287; Ipsilateral: F_(1,39)_=5.57, **p=.023**, η_p_=0.125). The ipsilateral interaction between group and difficulty was driven by an increase of activity with difficulty in the amputees (t_(39)_= 3.55, **p<.001**), but not in one-handers or controls (t_(39)_=-0.18, p=.857; t_(39)_= 0.90, p=. 373). Furthermore, amputees showed significantly larger activity than controls in the ipsilateral cortex for both the difficult (t_(39)_= 3.37, **p=.002**) and the easy conditions (t_(39)_= 3.22, **p=.003**). Together, these findings confirm and extend previous studies – ipsilateral activity for the intact hand was heightened in amputees, particularly with increased task difficulty. Conversely, the one-handed group did not show any significant benefit or disadvantage in activating the missing hand cortex relative to controls.

**Figure 2.**
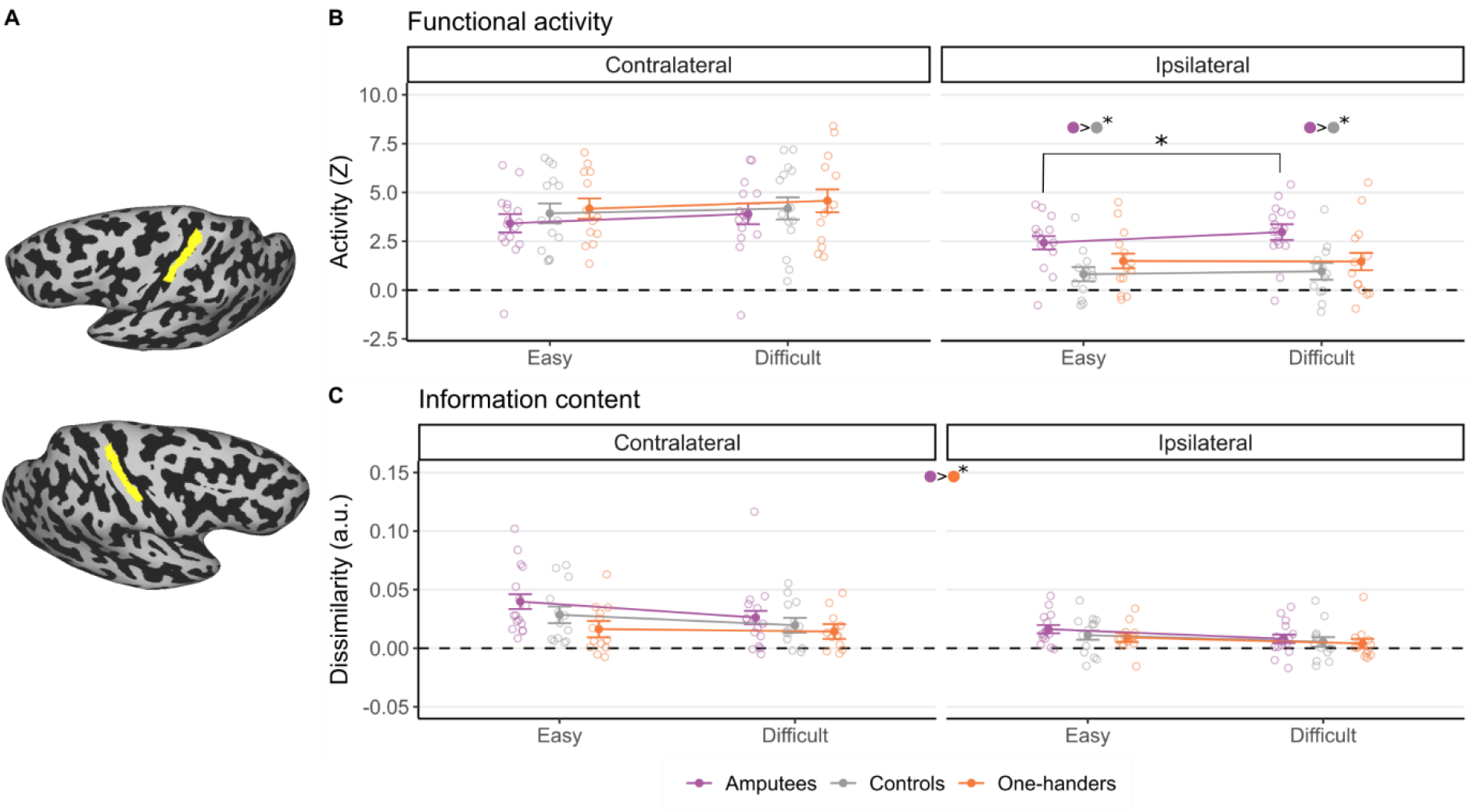
Bilateral univariate and multivariate fMRI activity of the intact hand for easy and difficult multi-finger configurations. **(A)** Bilateral somatosensory (BA3b) hand ROIs used in the analyses (one example participant). **(B)** Brain activity (Z-scores, calculated independently for each configuration versus rest) in the contralateral and ipsilateral hemispheres, averaged for the easy (145 and 234) and difficult (134, 125, and 235) configurations across runs. **(C)** Information content (dissimilarities between configuration pairs) in the contralateral and ipsilateral hemispheres averaged across runs. Only the dissimilarities between the easiest (e.g., the green square in Figure 3A) and most difficult (e.g., the blue square in Figure 3A) finger configurations were selected. The open dots represent individual participants. Colour filled dots at the top of plots B and C indicate significant difference between the groups for the most relevant comparisons. Statistical significance (Bonferroni corrected) is indicated as follows: *: significance difference; #: trending difference (p < 0.07). Amputees showed significantly larger activity than the control and one-hander groups in the ipsilateral cortex for the difficulty condition, however this did not result in an increased information content **BA3b ROI.** Amputees showed significantly larger activity than the control and one-hander groups in the ipsilateral cortex for the difficulty condition (Panel B) but this did not result in an increased information content (Panel C). **A)** Bilateral hand BA3b ROIs used in the analyses (one example participant). **B)** Brain activity (zscores) in the contralateral and ipsilateral hemispheres averaged across runs and across easy and difficult configurations. **C)** Information content (dissimilarities between configuration pairs) in the contralateral and ipsilateral hemispheres averaged across runs. Only the dissimilarities between the easiest (e.g., the green square in Figure 3A) and most difficult (e.g., the blue square in Figure 3A) finger configurations were selected. The unshaded dots with different colours represent individual participants. Colour filled dots with asterisks at the top of plots B and C indicate significant difference (Bonferroni corrected) between the groups specified by the colours in a specific or averaged condition. Lines with asterisks refer to significant difference (Bonferroni corrected) between conditions within a group. We are reporting here only the relevant comparisons, for the complete analysis, please refer to the results section.

We repeated the same analysis in bilateral M1 hand ROI. The 3-way interaction was not significant in this case (F_(2,39)_=1.50, p=.236, η_p_=0.07). We observed a significant interaction between difficulty and hemisphere (F_(1,39)_=11.48, **p=.002**, η_p_=0.23), driven by activity increase with difficulty in the contralateral hemisphere only (Contralateral: t_(39)_=4.07, p=.0002; Ipsilateral: t_(39)_=2.02, p=.05, Bonferroni adjusted α: .05/2=.025). We also observed as a significant interaction between group and hemisphere (F_(2,39)_=4.83, **p=.013**, η_p_=0.20), due to the fact that the difference in activity between the two hemispheres was reduced in the amputees relative to the control groups (t_(39)_= −3.05, p=0.004, Bonferroni adjusted α: .05/3=.0167). This is in line with the observation that the amputees showed higher activity in the ipsilateral hemisphere than the control group. We did not find main effects or interaction between group and difficulty (all p>.3). Overall, these results suggest that the Finally, to confirm that our effects reflect increased difficulty relating to motor performance per se, rather than more general task demands, e.g. relating to attentional or arousal effects, we repeated the same analysis in a control visual region (left hMT+) and observed no significant main effects or interactions (all p>.15).

### Amputees show bilateral increase in information content relative to one-handers

We next assessed whether the selective increase in unilateral activity observed in amputees, previously interpreted as functional remapping, translated to a gain in information content. Average distances (across all configuration pairs as shown in Figure 3A) were significantly greater than zero (all ps<.05, not corrected for multiple comparisons), confirming that task relevant information was encoded in both hemispheres. We first examined distances when specifically comparing the easy and difficult configurations separately across hemispheres and groups. To allow us to specifically account for difficulty, this analysis was restricted to the easiest configuration pair (C234-C145) and the most difficult configuration pair (C235-C125) in our representational dissimilarity matrix (highlighted in Figure 3A, green: easiest; blue: most difficult). If increase in activity translate to information content gain, we should see larger distances between configuration pairs for the amputees, especially across the most difficult conditions. However, we did not find a significant 3-way interaction (F_(2,38)_=0.41, p=.668, η_p_=0.15), or a resulting 2-way interaction involving group (see Figure 2C). Instead, we found a main effect of group (F_(2,38)_=3.29, **p=.048**, η_p_=0.15) driven by increased information content in amputees relative to one-handers (t_(38)_= 2.55, **p=.015**, Bonferroni adjusted α: .05/3=.0167). We also observed a main effect of hemisphere (F_(1,38)_=15.57, **p=<.001**, η_p_=0.29) and difficulty (F_(1,38)_=11.91, **p=.001**, η_p_=0.24). Interestingly, we found that information scales down with difficulty, regardless of group, suggesting that the overall increase in information observed in amputees is not linked to their reduced performance.

**Figure 3.**
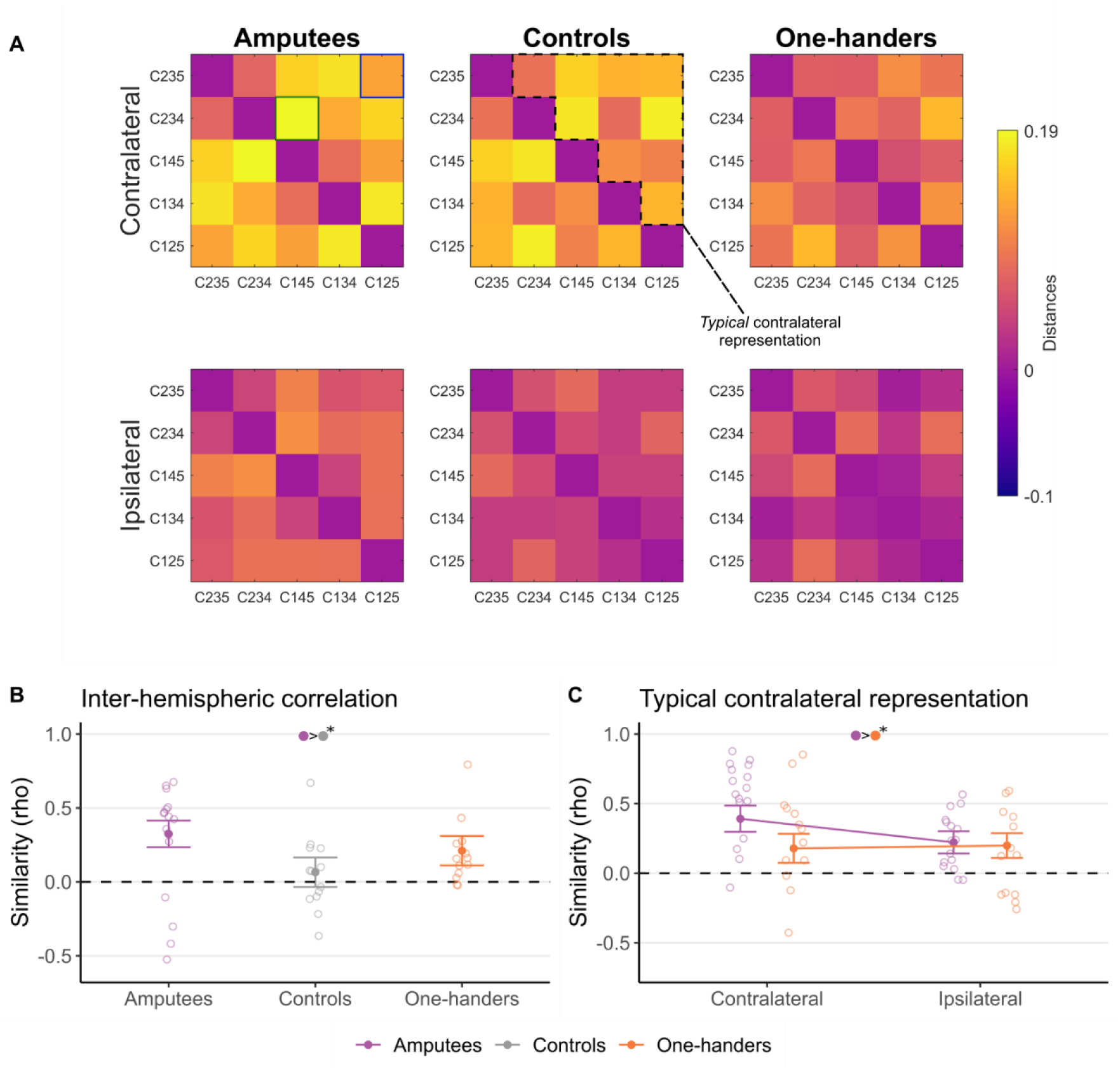
Functional homotopy and contralateral typicality in multivariate representational structure. **A)** Representation dissimilarity matrices (RDMs) across multi-finger configurations, groups and hemispheres. Colours reflect crossnobis distance, with warmer distances showing greater pairwise dissimilarity. The green and blue squares on the top left RDM highlight the easiest (green) and most difficult (blue) finger configuration pairs, respectively for the analysis in Figure 2C (the same pairs were used for the other RDMs). The dashed area on the Contralateral Controls RDM indicate the typical contralateral representation used to assess the typicality of representation in Figure 3C. **B)** Inter-hemispheric correlation (rho) between the contralateral and ipsilateral RDM within individuals was used calculate homotopy. **C)** The individual RDMs of the amputees and one-handers groups were correlated with the average contralateral RDM of the controls (the typical contralateral representation) to calculate contra-typical representation. All other annotations are as reported in Figure legend 2. Amputees showed typical contralateral representational motifs in their missing hand cortex for representing multi-finger configurations with their intact hand.

To take best advantage of our information content analysis, we repeated the analysis while comparing the average distances across the entire RDM (10 cells) across groups and hemispheres in a 2-way ANCOVA. Again, if increase in activity translate to information content gain, we should see larger averaged distance between configuration pairs for the amputees. Here again, we find no significant interaction (F_(2,38)_=0.65, p=.525, η_p_=0.03), suggesting that information content was not modulated differently across group and hemisphere. Instead, again, we find a main effect of group (F_(2,38)_=3.83, **p=.030**, η_p_=0.17) and hemisphere (F_(1,38)_=25.84, **p<.001**, η_p_=0.40). The main effect of group was again driven by increased distances across both hemispheres in amputees relative to one-handers (t_(38)_= 2.62, **p=.013**). Similar to the previous analysis, these effects were not specific to the ipsilateral cortex, but were instead generalised. Do these group differences reflect increased information in amputees or decreased information in one-handers? When comparing against controls, the results are ambiguous (amputees versus controls: t_(38)_= 1.99, p=.054, BF_10_=0.90; one-handers versus controls: t_(38)_= −0.57, p=.573, BF_10_=0.46; Bonferroni adjusted α: 0.05/2=.025). Together, it appears that the increased activity found in the ipsilateral hemisphere of amputees for the difficult configurations does not neatly translate to a selective increased information content.

To further confirm the specificity of our effects, we repeated the same analyses in M1 and hMT+, and verified that the averaged distances were also significantly larger than zero (all p<.007, not corrected for multiple comparisons). In M1, when focusing on difficulty as a factor, we observed a main effect of difficulty (F_(1,38)_=5.60, **p=.023**, η_p_=0.13), suggesting larger distances between the easiest pairs than the most difficult ones, and hemisphere (F_(1,38)_=15.83, **p<.01**, η_p_=0.29), suggesting larges distances in the contralateral hemisphere relative to the ipsilateral hemisphere. No main effect of group (F_(2,38)_=2.05, p<.143, η_p_=0.10), and no group interactions. Similarly, when averaging across all configurations, we observed a main effect of hemisphere (F_(1,38)_=14.79, **p=<.001**, η_p_=0.28), suggesting larger distances in the contralateral relative to the ipsilateral hesmiphere, no main effect of group (F_(2,38)_=1.94, p=.158, η_p_=0.09), and no group interactions. Despite higher distances in hMT+ (presumably due to the visual information provided throughout the motor task), we did not observe any main effects or interactions (all p> 0.2).

### Amputees show increased functional homotopy in representational structure across hemispheres

Functional homotopy refers to brain regions in opposite hemispheres exhibiting correlated activity patterns during a task or at rest, and suggests that two brain regions are functionally associated and working in concert to perform a certain function (e.g., a motor task). We explored the degree of functional homotopy (defined here as the correlation between representational dissimilarity matrices shown in Figure 3A) in the hand region across the two hemispheres. We first correlated the 10 configuration pairs of the RDM across the two S1 hand areas of each participant. The homotopy correlation values were significantly larger than zero for the amputee (t_(15)_=2.83, **p=.013**) and one-hander groups (t_(12)_=3.28, **p=.006**), but not for the controls (t_(12)_=-0.59 **p=.563**, BF=0.32; Bonferroni adjusted α: .05/3=.016 for the 3 reported comparisons). When comparing across groups (using a 1-way ANCOVA, accounting for age), we found a trend towards significance (F_(2,38)_=2.99, p=.062, η_p_=.14, BF_10_=2.05), which is also reflected in greater homotopy in amputees relative to controls (t_(38)_=2.41, **p=.021**), but not for one-handers relative to controls (t_(38)_=1.63, p=.112, BF_10_=1.44; Bonferroni adjusted α: .05/2=.025 for the last 2 comparisons).

To determine whether the increased homotopy found in amputees reflects typical contralateral representation of the ipsilateral (missing hand) cortex, we next compared the ipsilateral representational structure of amputees and one-handers to the average RDM of controls’ contralateral average RDM. As shown in Figure 3C, for amputees we found a significant (above zero) correlation between both contralateral and ipsilateral ROIs relative to the typical contralateral representational structure in controls (Amputees Contralateral: t_(15)_=6.82, **p<.001;** Amputees Ipsilateral: t_(15)_= 4.38, **p<.001**, one-sample t-test; Bonferroni adjusted α: 0.05/2=0.025), whereas the correlation between one-handers and controls was approaching significance for the contralateral ROI only (One-handers contralateral: t_(12)_=2.51, **p=.027;** One-handers ipsilateral: t_(12)_=1.70, p=.114; Bonferroni adjusted α: 0.05/2=0.025). The two-way ANCOVA comparing group and hemisphere showed an expected effect of hemisphere (F_(1,26)_=9.44, **p=.005**, η_p_=0.26), reflecting the greater correlation of the contralateral hemisphere, and a significant main effect of group (F_(1,26)_= 4.64, **p=.041**, η_p_=0.15). The interaction was not significant (F_(1,26)_=1.73, p=.20, η_p_=0.06). This demonstrates that amputees represented the different finger configurations bilaterally in way that was similar to the typical representation in the contra-lateral hemisphere in neuro-typical controls.

When repeating the same set of analyses in M1, amputees only showed a significant correlation between the contralateral ROI relative to the typical contralateral structure in controls (Amputees Contralateral: t_(15)_=2.81, **p=.013;** Amputees Ipsilateral: t_(15)_= 0.45, p=.659; Bonferroni adjusted α: 0.05/2=0.025; One-handers Contralateral: t_(12)_= 1.62, p=.131; One-handers Ipsilateral: t_(12)_= 0.57, p=.575; Bonferroni adjusted α: 0.05/2=0.025). Furthermore, the two-way ANCOVA revealed a significant main effect of hemisphere (F_(1,26)_=5.55, **p=.026**, η_p_=0.17), no main effect of group (F_(1,26)_= 0.04, p=.840, η_p_=0.02) and no interaction (F_(1,26)_=0. 61, p=.443, η_p_=.02).

### No differences in white matter tracts between the three groups

Finally, we analysed diffusion MRI data, collected in the same cohort, to explore whether the group differences observed in the functional analysis are also reflected by alterations in structural connectivity. As noted in the Introduction, it is possible that ipsilateral functionality depends on the brain establishing (through bimanual experience) a functional interaction between the two hand territories. One possibility is that this is mediated, at least in part, via transcallosal pathways that connect the two hand areas (Fling et al., 2013). To address this, we conducted deterministic tractography to examine potential differences in the tissue microstructural properties of the transcallosal fibers connecting the two hand areas. We first compared the vertex-weighted mean FA and MD, derived from tractography-based inter-hemispheric connections, using two separate ANCOVAs (controlling for age). For both metrics, the main effect of group was not significant (FA: F_(2,35)_= 0.05, p=<.950, η_p_=.003, BF_10_=0.19; MD: F_(2,35)_= 0. 08, p=<.922, η_p_=.005, BF_10_=0.20). The Bayes

Factors in both analyses provided evidence in favour of the null hypothesis being no group structural differences in FA and MD.

To explore potential differences between amputees/one-handers and controls beyond these transcallosal interhemispheric connections, we conducted a complementary voxel-wise TBSS analyses at the whole brain level, as well as within a corticospinal tract ROI (see Methods; Hahamy et al., 2015). At the whole brain level, we found no FA or MD differences between either group (amputees and one-handers) or controls (TFCE-corrected, p=0.05). We also saw no significant clusters when contrasting amputees with one-handers. We did, however, find a negative effect of age, confirming the quality of the data. For the corticospinal tract, we similarly found no significant differences between each experimental group and the controls (both FA and MD), and this was also the case when comparing amputees with one-handers. Together, these findings do not support substantial structural changes in white matter architecture most relevant for inter-hemispheric coordination for motor control in our experimental groups.

## Discussion

In this study, we investigated the effect of missing a hand, whether it be due to congenital malformation or acquired amputation, on motor ability and representation of the intact hand. Given the profound behavioural pressure of growing up and/or living with only one hand, perceptual learning combined with practice effects are likely to enhance motor skills of the intact hand in both groups. Due to a combination of physiological and cognitive reasons, critical periods in development may be more favourable for training effects to occur (Sur and Rubenstein, 2005; Levelt and Ḧubener, 2012), making one-handers the most likely candidates to benefit from brain plasticity to improve motor control and learning with the intact hand. Instead, we found that one-handers showed poorer performance in a finger configuration task, particularly with regards to motor learning of the more difficult configurations. In contrast, amputees did not show any clear deviations from controls during task training outside the scanner. This finding aligns with prior research indicating motor deficits in one-handers but not amputees. For example, one-handers (Philip et al., 2015) but not amputees (Philip and Frey, 2011) exhibited accuracy and speed deficits while planning a grasp with their intact hand. Based on this, it has been postulated that sensorimotor experience of both hands is necessary for the refinement of accurate unilateral motor prediction and performance (Philip et al., 2015). Relatedly, we previously found that one-handers made more errors during visually guided reaching with their artificial arm, relative to amputees, as well as two-handed controls using their nondominant arm (Maimon-Mor et al., 2021) (though it is important to point that in this study intact hand reaching performance was not significantly different from the other groups). Interestingly, one-handers who started using an artificial arm for the first time earlier as toddlers showed less motor deficit, hinting at a critical period for integrating a visuomotor representation of a limb. Together, these findings imply that a disability experienced in early life may impede motor development, even for body parts not directly affected by the malformation. This reasoning does not necessarily contradict the more straightforward prediction that motor control and learning would be superior in one-handers due to early life behavioural pressure. It is possible that critical periods trigger both long-term deficits and improved skill that would counterbalance each other. According to this rationale, if it wasn’t for over-practice in early life, one-handers would show more severe motor impairments in their daily life.

Despite heightened activity in the territory of the missing hand during task performance (as discussed below), amputees did not display superior motor performance with their intact hand. The idea that amputees develop enhanced behavioural abilities following their amputation due to reallocation of central resources in the missing hand cortex has been a topic of much fascination for the past century. Originally, hypotheses (and reports) focused on heightened tactile sensitivity on the residual limb (stump) of human amputees (e.g., Katz, 1920; Teuber, H and Krieger, HP and Bender, 1949; Haber, 1955) (see Makin, 2021 for maladaptive consequences of reorganisation in amputees). Merzenich and colleagues (1984) proposed that remapping following finger amputation should lead to increased tactile acuity of the neighbouring fingers. Other studies, using short-term and reversible deafferentation, suggested that, due to increased excitability of the deafferented hemisphere, amputation should result in increased acuity for the non-deafferented (‘intact’) hand (Björkman et al., 2004; Lissek et al., 2009; Dempsey-Jones et al., 2019). More recently, we and other suggested that increased activity for the intact hand in the missing hand territory is a potential neural correlate of adaptive plasticity for motor abilities (Makin et al., 2013a; Philip and Frey, 2014). The assumption behind these ideas is that the brain can correctly interpret signals arising from the missing hand territory as relating to the intact hand, thereby providing greater (or better) information about the new representation. This is consistent with physiological studies showing that, while hand and finger movements are mostly controlled through crossed corticospinal projections from the contralateral hemisphere (Brinkman and Kuypers, 1973), there are also known ipsilateral (uncrossed) motor projections (Soteropoulos et al., 2011). Given that amputees are pressured to rely on their remaining hand heavily in their day-to-day activities, one might expect improved read-out of neural signals originating from the ipsilateral cortex, which typically has limited functionality in individuals with both hands intact. This improvement should lead to recruitment of the missing hand hemisphere in the brain’s ipsilateral region. However, much of the original evidence for perceptual gains in amputees have been since challenged (O’Boyle et al., 2001; Vega-Bermudez and Johnson, 2002) In our brief training paradigm, we found no evidence for motor behavioural benefits in amputees.

What is then the functional relevance of the increased ipsilateral activity observed in sensorimotor cortex of amputees here, as well as in many previous studies (Kew et al., 1994; Hamzei et al., 2001; Bogdanov et al., 2012; Makin et al., 2013a; Philip and Frey, 2014; Valyear et al., 2020)? One difficulty in interpreting the functional meaning of net changes in activity levels is that they could result from multiple dissociated mechanisms, such as aberrant processing (Makin et al., 2013b), disinhibition (Hahamy et al., 2017), or merely reflect gain changes due to upstream processing (Kambi et al., 2014). Common to these alternative processes is that increased activity doesn’t necessarily entail a change of the underlying information being processed (Arbuckle et al., 2019). In other words, activity changes that underlie remapping do not necessarily entail information content changes. In the present study, we found that, while difficulty increases contralateral activity across all groups, in the ipsilateral cortex difficulty increases activity significantly only in amputees. This is interesting, because it goes against the idea that the increased ipsilateral activity is a simple passive consequence of inter-hemispheric disinhibition (Werhahn et al., 2002; Ramachandran and Altschuler, 2009; Simões et al., 2012). Instead, it seems that the ipsilateral cortex is selectively recruited for the more difficult configurations in amputees. However, this result should not be taken as evidence for functional recruitment of ipsilateral cortex. Representational similarity analysis (RSA) is a multivariate technique designed to determine how separate or distinct one activity pattern is to another. RSA allows us to ask not only if more information is available in a given brain area (dissimilarity distances), but also whether this new information is structured consistently with known representational principles, e.g. related to the contralateral hemisphere. By quantifying and characterising brain function beyond the spatial attributes of activity maps, while providing a more precise model for how information content varies across configurations, we believe RSA provides an arguably better tool for assessing the functional characteristics of the ipsilateral cortex. Furthermore, both in our previous study (conducted on the same set of participants Wesselink et al., 2019) and here, we found that the increased activity in amputees did not translate to differentially increased ipsilateral information.

Several previous studies using multivariate pattern analysis have demonstrated that, despite activity suppression, ipsilateral sensory and motor cortex contains information pertaining to individual fingers (Diedrichsen et al., 2013b, 2018; Berlot et al., 2019; Wesselink et al., 2019). These ipsilateral activity patterns appear to be weaker, but otherwise similar in representational structure to those elicited by movement of the mirror-symmetric finger in the opposing hand, at least for single finger movements (Diedrichsen et al., 2013a, 2018). The ipsilateral representation is not a simple ‘spill-over’ or passive copy from the homologous (contralateral) hand area, as it has been shown to be differently modulated by behavioural task context (Berlot et al., 2019). Yet, the functional significance of these ipsilateral representations and independence from the contralateral representation is still unknown. Ipsilateral activity in M1 has also been observed in monkey studies during proximal (i.e., shoulders and elbows) motor tasks (Ames and Churchland, 2019; Heming et al., 2019; Cross et al., 2020). These studies seem to suggest that, even if the same population of neurons encodes both ipsilateral and contralateral movements, the two limb representations are distributed differently across neurons (i.e., arm-related activity occupies distinct subspaces), which is proposed to be the mechanism that avoid impacting (i.e., moving) the wrong arm. Furthermore, it has been suggested that the ipsilateral representation is an *efference* copy resulting from the contralateral activity to inform the ipsilateral cortex about the contralateral arm movement and help with bimanual coordination (Ames and Churchland, 2019). The efference copy would be sent by default, even in the absence of bimanual movements, and ignored if not needed. In other words, the ipsilateral representation could be a consequences of the fact that the two homologous areas inform each other about their respective current state. Although the relationship between level of task complexity and the functional role of the efference copy has not been explored yet, it is interesting to speculate that the relevance of the efference copy will be greater for tasks requiring coordination across hands.

This latter interpretation provides an interesting conceptual framework for our reported findings: in one-handers, the lack of bimanual experience will dampen the mechanistic development of bimanual hand representation, including cross-hemisphere efference copy, resulting in bilateral reduction of information content. Whereas in amputees – the ipsilateral efference copy from the intact hand will be more prominent in the missing hand cortex due to the reduced utilisation of the missing hand, resulting in increased homotopy for the intact hand across the two hemispheres. Importantly, under this conceptual framework, we should not expect that representational changes will have a functional behavioural impact on our participants. This is because the efference copy that is presumably being modulated is designed to improve bimanual coordination, which is impossible for amputees to implement. This interpretation is consistent with a recent study which did not find a functional relevance for increased S1 ipsilateral activity (Valyear et al., 2020), in line with our observation that amputees did not show any behavioural improvement outside the scanner and, if anything, they showed a performance reduction inside the scanner. Our white matter findings also provide indirect support for the functional irrelevance of activity changes, as it provides substantial evidence to support an account of stable anatomy despite increased activity and better inter-hemispheric collaboration. In this context, it is interesting to consider previous evidence for persistent representation of the missing hand in amputees despite amputation (Raffin et al., 2012; Makin et al., 2013b). It is interesting to speculate whether the homotopic representation of the intact hand in the missing hand cortex helps maintain the missing hand representation. While we and others previously showed that the phantom hand map is activated by phantom hand movements independently of the intact hand (Kikkert et al., 2016; Bruurmijn et al., 2017; Wesselink et al., 2019), it is still possible that structured inputs from the intact hand (via ipsilateral pathways) sustains the missing hand map, despite the loss of the original peripheral inputs.

To conclude, our findings reveal a collaborative relationship between the contralateral and ipsilateral cortices during task performance in amputees, above and beyond the other groups. By focusing on information content and its representational structure above and beyond the salient effects of remapping, defined as increased mean activity, our findings highlight a different aspect of the critical period than normally emphasised, which is based on experience rather than deprivation. Specifically, representations of both hands and some bimanual experience in the early developmental stage is necessary to develop a bilateral motor representation and a typical contralateral representation. Interestingly, while the ipsilateral efference copy interpretation is functionally irrelevant for the unimanual tasks studied here and in previous research, it may provide a useful consideration, and perhaps even exciting new opportunities, for combining novel restorative brain-computer interfaces (Fouad et al., 2015) and augmentation technologies (Dominijanni et al., 2021) for bimanual interactions.

## References

Ames KC, Churchland MM (2019) Motor cortex signals for each arm are mixed across hemispheres and neurons yet partitioned within the population response. Elife 8:1–36.

Amunts K, Mohlberg H, Bludau S, Zilles K (2020) Julich-Brain: A 3D probabilistic atlas of the human brain’s cytoarchitecture. Science (80-) 369:988–992.

Andersson JLR, Jenkinson M, Smith S (2010) Non-linear registration, aka spatial normalization (FMRIB technical report TR07JA2). FMRIB Anal Gr Univ Oxford.

Andersson JLR, Skare S, Ashburner J (2003) How to correct susceptibility distortions in spin-echo echo-planar images: Application to diffusion tensor imaging. Neuroimage 20:870–888.

Arbuckle SA, Pruszynski JA, Diedrichsen J (2022) Mapping the Integration of Sensory Information across Fingers in Human Sensorimotor Cortex. J Neurosci 42:5173–5185 Available at: https://www.jneurosci.org/lookup/doi/10.1523/JNEUROSCI.2152-21.2022.

Arbuckle SA, Yokoi A, Pruszynski JA, Diedrichsen J (2019) Stability of representational geometry across a wide range of fMRI activity levels. Neuroimage 186:155–163.

Berlot E, Prichard G, O’Reilly J, Ejaz N, Diedrichsen J (2019) Ipsilateral finger representations in the sensorimotor cortex are driven by active movement processes, not passive sensory input. J Neurophysiol 121:418–426.

Björkman A, Rosén B, Van Westen D, Larsson EM, Lundborg G (2004) Acute improvement of contralateral hand function after deafferentation. Neuroreport 15:1861–1865.

Bogdanov S, Smith J, Frey SH (2012) Former hand territory activity increases after amputation during intact hand movements, but is unaffected by illusory visual feedback. Neurorehabil Neural Repair 26:604–615.

Brinkman J, Kuypers HGJM (1973) CEREBRAL CONTROL OF CONTRALATERAL AND IPSILATERAL ARM, HAND AND FINGER MOVEMENTS IN THE SPLIT-BRAIN RHESUS MONKEY. Brain 96:653–674 Available at: https://academic.oup.com/brain/article-lookup/doi/10.1093/brain/96.4.653.

Bruurmijn MLCM, Pereboom IPL, Vansteensel MJ, Raemaekers MAH, Ramsey NF (2017) Preservation of hand movement representation in the sensorimotor areas of amputees. Brain 140:3166– 3178.

Chang LC, Jones DK, Pierpaoli C (2005) RESTORE: Robust estimation of tensors by outlier rejection. Magn Reson Med 53:1088–1095.

Cross KP, Heming EA, Cook DJ, Scott SH (2020) Maintained Representations of the Ipsilateral and Contralateral Limbs during Bimanual Control in Primary Motor Cortex. J Neurosci 40:6732–6747 Available at: https://www.jneurosci.org/lookup/doi/10.1523/JNEUROSCI.0730-20.2020.

Dale AM, Fischl B, Sereno MI (1999) Cortical Surface-Based Analysis: I. Segmentation and Surface Reconstruction. Neuroimage 9:179–194 Available at: https://linkinghub.elsevier.com/retrieve/pii/S1053811998903950.

Dell’Acqua F, Tournier JD (2019) Modelling white matter with spherical deconvolution: How and why? NMR Biomed 32:1–18.

Dempsey-Jones H, Themistocleous AC, Carone D, Ng TWC, Harrar V, Makin TR (2019) Blocking tactile input to one finger using anaesthetic enhances touch perception and learning in other fingers. J Exp Psychol Gen 148:713–727 Available at: http://doi.apa.org/getdoi.cfm?doi=10.1037/xge0000514.

Diedrichsen J, Provost S, Zareamoghaddam H (2016) On the distribution of cross-validated Mahalanobis distances. arXiv Prepr:1–24 Available at: http://arxiv.org/abs/1607.01371.

Diedrichsen J, Wiestler T, Ejaz N (2013a) A multivariate method to determine the dimensionality of neural representation from population activity. Neuroimage 76:225–235 Available at: http://dx.doi.org/10.1016/j.neuroimage.2013.02.062.

Diedrichsen J, Wiestler T, Krakauer JW (2013b) Two distinct ipsilateral cortical representations for individuated finger movements. Cereb Cortex 23:1362–1377.

Diedrichsen J, Yokoi A, Arbuckle SA (2018) Pattern component modeling: A flexible approach for understanding the representational structure of brain activity patterns. Neuroimage 180:119– 133 Available at: https://linkinghub.elsevier.com/retrieve/pii/S1053811917306985.

Dienes Z (2014) Using Bayes to get the most out of non-significant results. Front Psychol 5:1–17.

Dominijanni G, Shokur S, Salvietti G, Buehler S, Palmerini E, Rossi S, De Vignemont F, D’Avella A, Makin TR, Prattichizzo D, Micera S (2021) The neural resource allocation problem when enhancing human bodies with extra robotic limbs. Nat Mach Intell 3:850–860.

Ejaz N, Hamada M, Diedrichsen J (2015) Hand use predicts the structure of representations in sensorimotor cortex. Nat Neurosci 18:1034–1040 Available at: http://www.nature.com/articles/nn.4038.

Fischl B, Liu a, Dale a M (2001) Automated manifold surgery: constructing geometrically accurate and topologically correct models of the human cerebral cortex. IEEE Trans Med Imaging 20:70–80 Available at: http://www.ncbi.nlm.nih.gov/pubmed/11293693.

Fischl B, Rajendran N, Busa E, Augustinack J, Hinds O, Yeo BTT, Mohlberg H, Amunts K, Zilles K (2008) Cortical folding patterns and predicting cytoarchitecture. Cereb Cortex 18:1973–1980.

Fling BW, Benson BL, Seidler RD (2013) Transcallosal sensorimotor fiber tract structure-function relationships. Hum Brain Mapp 34:384–395 Available at: https://onlinelibrary.wiley.com/doi/10.1002/hbm.21437.

Fouad MM, Amin KM, El-Bendary N, Hassanien AE (2015) Brain Computer Interface: A Review. In: Brain-Computer Interfaces (Hassanien AE, Azar AT, eds), pp 3–30 Intelligent Systems Reference Library. Cham: Springer International Publishing. Available at: https://link.springer.com/10.1007/978-3-319-10978-7_1.

Glasser MF, Coalson TS, Robinson EC, Hacker CD, Harwell J, Yacoub E, Ugurbil K, Andersson J, Beckmann CF, Jenkinson M, Smith SM, Van Essen DC (2016) A multi-modal parcellation of human cerebral cortex. Nature 536:171–178 Available at: http://www.nature.com/doifinder/10.1038/nature18933.

Graziano MSA, Aflalo TN (2007) Mapping behavioral repertoire onto the cortex. Neuron 56:239–251.

Haber WB (1955) Effects of Loss of Limb on Sensory Functions. J Psychol 40:115–123 Available at: http://www.tandfonline.com/doi/abs/10.1080/00223980.1955.9712969.

Hahamy A, Macdonald SN, van den Heiligenberg F, Kieliba P, Emir U, Malach R, Johansen-Berg H, Brugger P, Culham JC, Makin TR (2017) Representation of Multiple Body Parts in the Missing-Hand Territory of Congenital One-Handers. Curr Biol 27:1350–1355 Available at: http://dx.doi.org/10.1016/j.cub.2017.03.053.

Hahamy A, Makin TR (2019) Remapping in Cerebral and Cerebellar Cortices Is Not Restricted by Somatotopy. J Neurosci 39:9328–9342 Available at: https://www.jneurosci.org/lookup/doi/10.1523/JNEUROSCI.2599-18.2019.

Hahamy A, Sotiropoulos SN, Henderson Slater D, Malach R, Johansen-Berg H, Makin TR (2015) Normalisation of brain connectivity through compensatory behaviour, despite congenital hand absence. Elife 4:1–12 Available at: https://elifesciences.org/articles/04605.

Hamzei F, Liepert J, Dettmers C, Adler T, Kiebel S, Rijntjes M, Weiller C (2001) Structural and functional cortical abnormalities after upper limb amputation during childhood. Neuroreport 12:957–962.

Heming EA, Cross KP, Takei T, Cook DJ, Scott SH (2019) Independent representations of ipsilateral and contralateral limbs in primary motor cortex. Elife 8:1–26.

Jenkinson M, Bannister P, Brady M, Smith S (2002) Improved Optimization for the Robust and Accurate Linear Registration and Motion Correction of Brain Images. Neuroimage 17:825–841.

Jenkinson M, Beckmann CF, Behrens TEJ, Woolrich MW, Smith SM (2012) FSL. Neuroimage 62:782– 790 Available at: https://linkinghub.elsevier.com/retrieve/pii/S1053811911010603.

Kambi N, Halder P, Rajan R, Arora V, Chand P, Arora M, Jain N (2014) Large-scale reorganization of the somatosensory cortex following spinal cord injuries is due to brainstem plasticity. Nat Commun 5:3602 Available at: http://www.nature.com/articles/ncomms4602.

Katz D (1920) Psychologische versuche mit amputierten, Z. Psychol. Barth.

Kellner E, Dhital B, Kiselev VG, Reisert M (2016) Magnetic Resonance in Med - 2015 - Kellner - Gibbs-ringing artifact removal based on local subvoxel-shifts.pdf. Magn Reson Med 76:1574–1581.

Kew JJM, Ridding MC, Rothwell JC, Passingham RE, Leigh PN, Sooriakumaran S, Frackowiak RSJ, Brooks DJ (1994) Reorganization of cortical blood flow and transcranial magnetic stimulation maps in human subjects after upper limb amputation. J Neurophysiol 72:2517–2524.

Kikkert S, Kolasinski J, Jbabdi S, Tracey I, Beckmann CF, Berg HJ, Makin TR (2016) Revealing the neural fingerprints of a missing hand. Elife 5:1–19.

Kriegeskorte N, Mur M, Bandettini P (2008) Representational similarity analysis - connecting the branches of systems neuroscience. Front Syst Neurosci 2:4 Available at: http://www.pubmedcentral.nih.gov/articlerender.fcgi?artid=2605405&tool=pmcentrez&rendertype=abstract [Accessed July 13, 2012].

Leemans A, Jeurissen B, Sijbers J (2009) ExploreDTI: a graphical toolbox for processing.: 5–6.

Levelt CN, Ḧubener M (2012) Critical-period plasticity in the visual cortex. Annu Rev Neurosci 35:309– 330.

Lissek S, Wilimzig C, Stude P, Pleger B, Kalisch T, Maier C, Peters SA, Nicolas V, Tegenthoff M, Dinse HR (2009) Immobilization Impairs Tactile Perception and Shrinks Somatosensory Cortical Maps. Curr Biol 19:837–842.

Maimon-Mor RO, Schone HR, Slater DH, Faisal AA, Makin TR (2021) Early life experience sets hard limits on motor learning as evidenced from artificial arm use. Elife 10:1–26.

Makin TR (2021) Phantom limb pain: Thinking outside the (mirror) box. Brain 144:1929–1932.

Makin TR, Cramer AO, Scholz J, Hahamy A, Slater DH, Tracey I, Johansen-Berg H (2013a) Deprivation-related and use-dependent plasticity go hand in hand. Elife 2:1–15.

Makin TR, Flor H (2020) Brain (re)organisation following amputation: Implications for phantom limb pain. Neuroimage 218:116943 Available at: https://doi.org/10.1016/j.neuroimage.2020.116943.

Makin TR, Scholz J, Filippini N, Henderson Slater D, Tracey I, Johansen-Berg H (2013b) Phantom pain is associated with preserved structure and function in the former hand area. Nat Commun 4:1570–1578 Available at: http://dx.doi.org/10.1038/ncomms2571.

Merzenich MM, Nelson RJ, Stryker MP, Cynader MS, Schoppmann A, Zook JM (1984) Somatosensory cortical map changes following digit amputation in adult monkeys. J Comp Neurol 224:591–605 Available at: http://doi.wiley.com/10.1002/cne.902240408.

O’Boyle DJ, Moore CEG, Poliakoff E, Butterworth R, Sutton A, Cody FWJ (2001) Human locognosic acuity on the arm varies with explicit and implicit manipulations of attention: Implications for interpreting elevated tactile acuity on an amputation stump. Neurosci Lett 305:37–40.

Ogawa K, Mitsui K, Imai F, Nishida S (2019) Long-term training-dependent representation of individual finger movements in the primary motor cortex. Neuroimage 202:116051 Available at: http://www.ncbi.nlm.nih.gov/pubmed/31351164 [Accessed November 16, 2021].

Philip BA, Buckon C, Sienko S, Aiona M, Ross S, Frey SH (2015) Maturation and experience in action representation: Bilateral deficits in unilateral congenital amelia. Neuropsychologia 75:420–430.

Philip BA, Frey SH (2011) Preserved grip selection planning in chronic unilateral upper extremity amputees. Exp Brain Res 214:437–452.

Philip BA, Frey SH (2014) Compensatory Changes Accompanying Chronic Forced Use of the Nondominant Hand by Unilateral Amputees. J Neurosci 34:3622–3631 Available at: https://www.jneurosci.org/lookup/doi/10.1523/JNEUROSCI.3770-13.2014.

Postans M, Parker GD, Lundell H, Ptito M, Hamandi K, Gray WP, Aggleton JP, Dyrby TB, Jones DK, Winter M (2020) Uncovering a Role for the Dorsal Hippocampal Commissure in Recognition Memory. Cereb Cortex 30:1001–1015.

Pruszynski JA, Johansson RS, Flanagan JR (2016) A rapid tactile-motor reflex automatically guides reaching toward handheld objects. Curr Biol 26:788–792 Available at: http://dx.doi.org/10.1016/j.cub.2016.01.027.

R Core Team (2022) R: A language and environment for statistical computing. Accessed 1st April 2019 Available at: https://www.r-project.org/.

Raffin E, Mattout J, Reilly KT, Giraux P (2012) Disentangling motor execution from motor imagery with the phantom limb. Brain 135:582–595.

Raja Beharelle A, Dick AS, Josse G, Solodkin A, Huttenlocher PR, Levine SC, Small SL (2010) Left hemisphere regions are critical for language in the face of early left focal brain injury. Brain 133:1707–1716.

Ramachandran VS, Altschuler EL (2009) The use of visual feedback, in particular mirror visual feedback, in restoring brain function. Brain 132:1693–1710.

Schieber MH (2001) Constraints on Somatotopic Organization in the Primary Motor Cortex. J Neurophysiol 86:2125–2143 Available at: https://www.physiology.org/doi/10.1152/jn.2001.86.5.2125.

Schieber MH, Hibbard LS (1993) How somatotopic is the motor cortex hand area? Science (80-) 261:489–492.

Simões EL, Bramati I, Rodrigues E, Franzoi A, Moll J, Lent R, Tovar-Moll F (2012) Functional expansion of sensorimotor representation and structural reorganization of callosal connections in lower limb amputees. J Neurosci 32:3211–3220.

Smith SM, Jenkinson M, Johansen-Berg H, Rueckert D, Nichols TE, Mackay CE, Watkins KE, Ciccarelli O, Cader MZ, Matthews PM, Behrens TEJ (2006) Tract-based spatial statistics: Voxelwise analysis of multi-subject diffusion data. Neuroimage 31:1487–1505.

Smith SM, Jenkinson M, Woolrich MW, Beckmann CF, Behrens TEJ, Johansen-Berg H, Bannister PR, De Luca M, Drobnjak I, Flitney DE, Niazy RK, Saunders J, Vickers J, Zhang Y, De Stefano N, Brady JM, Matthews PM (2004) Advances in functional and structural MR image analysis and implementation as FSL. Neuroimage 23:208–219.

Soteropoulos DS, Edgley SA, Baker SN (2011) Lack of evidence for direct corticospinal contributions to control of the ipsilateral forelimb in monkey. J Neurosci 31:11208–11219.

Sur M, Rubenstein JLR (2005) Patterning and Plasticity of the Cerebral Cortex. Science (80-) 310:805– 810 Available at: https://onlinelibrary.wiley.com/doi/10.1002/9780470015902.a0000090.pub2.

Takesian AE, Hensch TK (2013) Balancing plasticity/stability across brain development. In: Progress in Brain Research, 1st ed., pp 3–34. Elsevier B.V. Available at: http://dx.doi.org/10.1016/B978-0-444-63327-9.00001-1.

Teuber, H and Krieger, HP and Bender M (1949) Reorganization of sensory function in amputation stumps: 2-point discrimination. Fed Proc 8:156--156.

Tillema JM, Byars AW, Jacola LM, Schapiro MB, Schmithorst VJ, Szaflarski JP, Holland SK (2008) Cortical reorganization of language functioning following perinatal left MCA stroke. Brain Lang 105:99– 111.

Tournier JD, Smith R, Raffelt D, Tabbara R, Dhollander T, Pietsch M, Christiaens D, Jeurissen B, Yeh CH, Connelly A (2019) MRtrix3: A fast, flexible and open software framework for medical image processing and visualisation. Neuroimage 202:116137 Available at: https://doi.org/10.1016/j.neuroimage.2019.116137.

Tuckute G, Paunov A, Kean H, Small H, Mineroff Z, Blank I, Fedorenko E (2022) Frontal language areas do not emerge in the absence of temporal language areas: A case study of an individual born without a left temporal lobe. Neuropsychologia 169:108184 Available at: https://doi.org/10.1016/j.neuropsychologia.2022.108184.

Valyear KF, Philip BA, Cirstea CM, Chen PW, Baune NA, Marchal N, Frey SH (2020) Interhemispheric transfer of post-amputation cortical plasticity within the human somatosensory cortex. Neuroimage 206:116291 Available at: https://doi.org/10.1016/j.neuroimage.2019.116291.

Vega-Bermudez F, Johnson KO (2002) Spatial acuity after digit amputation. Brain 125:1256–1264.

Veraart J, Novikov DS, Christiaens D, Ades-aron B, Sijbers J, Fieremans E (2016) Denoising of diffusion MRI using random matrix theory. Neuroimage 142:394–406 Available at: http://dx.doi.org/10.1016/j.neuroimage.2016.08.016.

Verstynen T, Diedrichsen J, Albert N, Aparicio P, Ivry RB (2005) Ipsilateral motor cortex activity during unimanual hand movements relates to task complexity. J Neurophysiol 93:1209–1222.

Vos SB, Tax CMW, Luijten PR, Ourselin S, Leemans A, Froeling M (2017) The importance of correcting for signal drift in diffusion MRI. Magn Reson Med 77:285–299.

Walther A, Nili H, Ejaz N, Alink A, Kriegeskorte N, Diedrichsen J (2016) Reliability of dissimilarity measures for multi-voxel pattern analysis. Neuroimage 137:188–200 Available at: http://dx.doi.org/10.1016/j.neuroimage.2015.12.012.

Waters-Metenier S, Husain M, Wiestler T, Diedrichsen J (2014) Bihemispheric transcranial direct current stimulation enhances effector-independent representations of motor synergy and sequence learning. J Neurosci 34:1037–1050.

Waters S, Wiestler T, Diedrichsen J (2017) Cooperation not competition: Bihemispheric tDCS and fMRI show role for ipsilateral hemisphere in motor learning. J Neurosci 37:7500–7512.

Werhahn KJ, Mortensen J, Kaelin-Lang A, Boroojerdi B, Cohen LG (2002) Cortical excitability changes induced by deafferentation of the contralateral hemisphere. Brain 125:1402–1413.

Wesselink DB, Heiligenberg FM Van Den, Ejaz N, Dempsey-Jones H, Cardinali L, Tarall-Jozwiak A, Diedrichsen J, Makin TR (2019) Obtaining and maintaining cortical hand representation as evidenced from acquired and congenital handlessness. Elife 8:1–19.

Wetzels R, Matzke D, Lee MD, Rouder JN, Iverson GJ, Wagenmakers EJ (2011) Statistical evidence in experimental psychology: An empirical comparison using 855 t tests. Perspect Psychol Sci 6:291– 298.

Wiestler T, Diedrichsen J (2013) Skill learning strengthens cortical representations of motor sequences. Elife 2:1–20 Available at: https://elifesciences.org/articles/00801.

Yousry TA, Schmid UD, Alkadhi H, Schmidt D, Peraud A, Buettner A, Winkler P (1997) Localization of the motor hand area to a knob on the precentral gyrus. A new landmark. Brain 120:141–157.

Yu WS, van Duinen H, Gandevia SC (2010) Limits to the Control of the Human Thumb and Fingers in Flexion and Extension. J Neurophysiol 103:278–289 Available at: https://www.physiology.org/doi/10.1152/jn.00797.2009.

